# CRP immunodeposition and proteomic analysis in abdominal aortic aneurysm

**DOI:** 10.1101/2020.12.30.424789

**Authors:** Eun Na Kim, Jiyoung Yu, Joon Seo Lim, Hwangkyo Jeong, Chong Jai Kim, Jae-Sung Choi, So Ra Kim, Hee-Sung Ahn, Kyunggon Kim, Se Jin Oh

**Author notes:** Computational Biology Program, Department of Biomedical Engineering, Oregon Health and Science University, Portland, OR, USA. Corresponding authors: (SJO), Email: (KK). These authors contributed equally to this work (Eun Na Kim, Jiyoung Yu). These authors also contributed equally to this work (Kyunggon Kim, Se Jin Oh).

## Abstract

**Objective:** The molecular mechanisms of the degeneration of the aortic wall in abdominal aortic aneurysm (AAA) are poorly understood. The monomeric form of C-reactive protein (mCRP) is deposited in damaged cardiovascular organs and aggravates the prognosis; however, it is unknown whether mCRP is deposited in the degenerated aorta of abdominal aortic aneurysm (AAA). We investigated whether mCRP is deposited in AAA and examined the associated pathogenic signaling pathways.

**Methods:** Twenty-four cases of AAA were analyzed and their histological features were compared according to the level of serum CRP and the degree of mCRP deposition. Proteomic analysis was performed in AAA cases with strong and diffuse CRP immunopositivity (n=7) and those with weak, focal, and junctional CRP immunopositivity (n=3).

**Results:** mCRP was deposited in the aortic specimens of AAA in a characteristic pattern that coincided with the lesion of the diminished elastic layer of the aortic wall. High serum CRP level was associated with stronger mCRP immunopositivity and a larger maximal diameter of aortic aneurysm. Proteomic analysis in AAA showed that multiple proteins were differentially expressed according to mCRP immunopositivity. Also, ingenuity pathway analysis showed that pathways associated with atherosclerosis, acute phase response, complement system, immune system, and coagulation were enriched in AAA cases with high mCRP immunopositivity.

**Conclusions:** AAA showed a characteristic deposition of mCRP, and multiple potentially pathologic signaling pathways were upregulated in AAA cases with strong CRP immunopositivity. mCRP and the aforementioned pathological pathways may serve as targets for managing the progression of AAA.

## INTRODUCTION

Abdominal aortic aneurysm (AAA), a permanent and progressive increase in the diameter of the aortic wall, is associated with 150,000–200,000 annual deaths worldwide[1]. Risk factors for AAA include old age, male sex, smoking, family history of AAA, hypertension, dyslipidemia, and the presence of other cardiovascular diseases such as ischemic heart disease or peripheral artery disease[2]. Human and animal studies showed that chronic aortic inflammation plays a major role in the pathogenesis of AAA[3]. However, the molecular mechanism of the degeneration of the aortic wall remains poorly understood, and an effective drug for preventing or inhibiting the degeneration has yet to be developed[4].

C-reactive protein (CRP) is an acute-phase protein that is rapidly produced from the liver and released into the bloodstream in response to various cellular injuries; as such, serum CRP level is widely used as a prognostic marker for cardiovascular diseases[5]. In terms of AAA, serum CRP level is considered as an independent risk factor[6], a predictor[7], and a prognostic marker[8]. Notably, CRP has different functions according to its structure. In the serum, CRP exists in a pentameric form (pCRP) and has an anti-inflammatory role[9]. However, when pCRP encounters damaged tissues, it dissociates into a monomeric form (mCRP) and is deposited in the damaged cell membrane[10]; subsequently, mCRP induces an inflammatory response by monocyte activation and reactive oxygen species formation to exacerbate tissue damage[11].

The deposition of mCRP has been reported in several cardiovascular diseases, including coronary artery atherosclerosis[12] and degenerated aortic valve[13]. Due to its role in tissue damage aggravation, the deposition of mCRP is associated with poor prognosis in cardiovascular diseases. However, whether mCRP is specifically deposited in the degenerated aorta is unknown. Here, we performed a thorough immunohistochemical evaluation on cases of AAA to determine whether mCRP is deposited in the damaged aortic walls, and if so, whether the degree of mCRP deposition is associated with proteomic changes thereof. We then compared the results of AAA with cases of ascending aortic dissection (AAD), which is largely derived from mechanical damage to the aorta and thus may serve as an alternative to healthy aorta samples for negative control without the biochemical effects of atherosclerosis. We also conducted quantitative proteomic analysis on the aorta specimens from AAA and AAD cases to examine the alterations in pathologic pathways in these conditions.

## MATERIALS AND METHODS

### Patient selection and clinical chart review

We collected cases of AAA and AAD that underwent aortic surgery at two tertiary referral centers in Seoul, Korea between January 2012 and December 2018. Among the AAA and AAD cases, we excluded those with inherited aortopathy or related connective tissue disorders such as Marfan syndrome, Loeys-Dietz syndrome, Ehlers-Danlos syndrome, Turner syndrome, and bicuspid aortic valve. Also, in order to exclude cases with elevations of serum CRP due to systemic causes, patients with acute systemic infections (e.g., influenza infection, pneumonia), inflammatory aortitis (e.g., IgG4-related disease, Takayasu aortitis, giant cell arteritis), or cancer aggravation were excluded. We also excluded patients in whom preoperative serum CRP level was not measured and those who had thoracic aortic aneurysm, thoracoabdominal aortic aneurysm, or combined aortic dissection with aortic aneurysm. Lastly, patients who previously underwent endovascular aortic surgery such as thoracic endovascular aortic repair or endovascular abdominal aortic aneurysm repair and those with iatrogenic or chronic dissection were also excluded.

A total of 69 patients with AAA and 87 patients with AAD were finally selected for analysis, and their clinical records were reviewed to retrieve the following data: age, sex, body mass index, medical history, presence of diabetes mellitus, hypertension, dyslipidemia, previous cardiovascular event including myocardial infarction, percutaneous coronary intervention, arteriosclerosis obliterans, medication history (e.g., statin), alcohol intake, smoking, preoperative serum CRP level, and serum WBC count. The largest aortic diameter in aortic aneurysm was analyzed by reviewing the preoperative aortic CT scan.

The study protocol was approved by the institutional review boards (IRBs) of Asan Medical Center (IRB number: S2020-0196) and SMG-SNU Boramae Medical Center (IRB number: 20190703/10-2019-54/081) and conformed to the relevant ethical guidelines and regulations. Because the bioethical law in South Korea mandates the need for written informed consent when using biological specimens obtained since 2013, we conducted the proteomic analysis using formalin-fixed paraffin-embedded (FFPE) tissue blocks obtained in 2012 with anonymized patient information. As such, both IRBs approved the use of clinical data as well as the collection and utilization of biological samples for research purposes and waived the need for formal written informed consent.

### Histopathologic analysis and immunohistochemical staining

During pathologic examinations of the surgical specimen, each aortic tissue was cut into 4mm thick sections after a detailed gross visual examination. Two or three cassettes were made by selecting the most severe and dilated portions of the aortic aneurysm accompanied by atherosclerosis (Supplementary Fig S1). The sectioned aorta specimens with atheroma were fixed in 10% buffered formalin and embedded in paraffin. The tissue sections were stained with hematoxylin and eosin (H&E).

Among the patients who underwent aortic surgery in 2012, 24 consecutive cases of AAA and six consecutive cases of AAD met the selection criteria, and the H&E slides of aortic wall specimen from those cases were histopathologically reviewed by a pathologist (E.N.K.). In AAA, the areas in which the thinning of the aortic wall was observed along with the atheromatous plaques were selected; in AAD, the areas in which medial tearing of the aortic wall was observed without atheromatous plaque were selected. Immunohistochemical staining was performed using the anti-CRP antibody (rabbit polyclonal, ab32412, Abcam, Cambridge, UK) that detects both the monomeric and pentameric forms of CRP, and the anti-mCRP antibody (mouse monoclonal, C1688, Sigma-Aldrich, Saint Louis, MO, USA) that selectively detects the monomeric form of CRP (24 kD subunit)[14, 15]. Masson trichrome special staining (Trichrome III Blue Staining Kit, Ventana Medical Systems, Tucson, AZ, USA) was performed to evaluate the degree of fibrosis with collagen deposition. Elastic staining (Elastic Staining Kit, Ventana Medical Systems) was performed to evaluate the elastic lamellae of aortic media and the structural changes in the aortic wall extracellular matrix to determine the medial degeneration of the aortic wall.

Additionally, we performed immunohistochemical staining with antibodies for monocyte chemoattractant protein-1 (MCP-1, ab9669, Abcam), CD68 for macrophages (M0814, Clone KP1, DAKO, Glostrup, Denmark), and complement markers including C1q (ab75756, Abcam), C3a (LS-B15388, LSBio, Seattle, USA), C5a (ab193295, Abcam), and C5a R (ab59390, Abcam). For secondary antibody, the Optiview DAB IHC Detection Kit (Ventana medical systems) was used. Immunopositive areas were classified according to the distribution of the immunopositivity of anti-CRP and anti-mCRP antibodies. Immunonegativity in both atheroma and junctional areas was classified as “negative”; none-to-weak immunopositivity in atheroma with strong, linear immunopositivity in junctional areas was classified as “junctional positive”; strong immunopositivity in both atheroma and junctional areas was classified as “diffuse positive” (Fig 1). Staining intensity was graded according to the atherosclerosis core, in which qualitative scores between 0 and 3 (negative [0], weak [1+], moderate [2+], strong [3+]) were applied (Fig 2). Pathologic evaluation and CRP immunoscoring of aortas were performed by a pathologist (E.N.K.) who was blinded to the clinical data.

**Fig 1.**
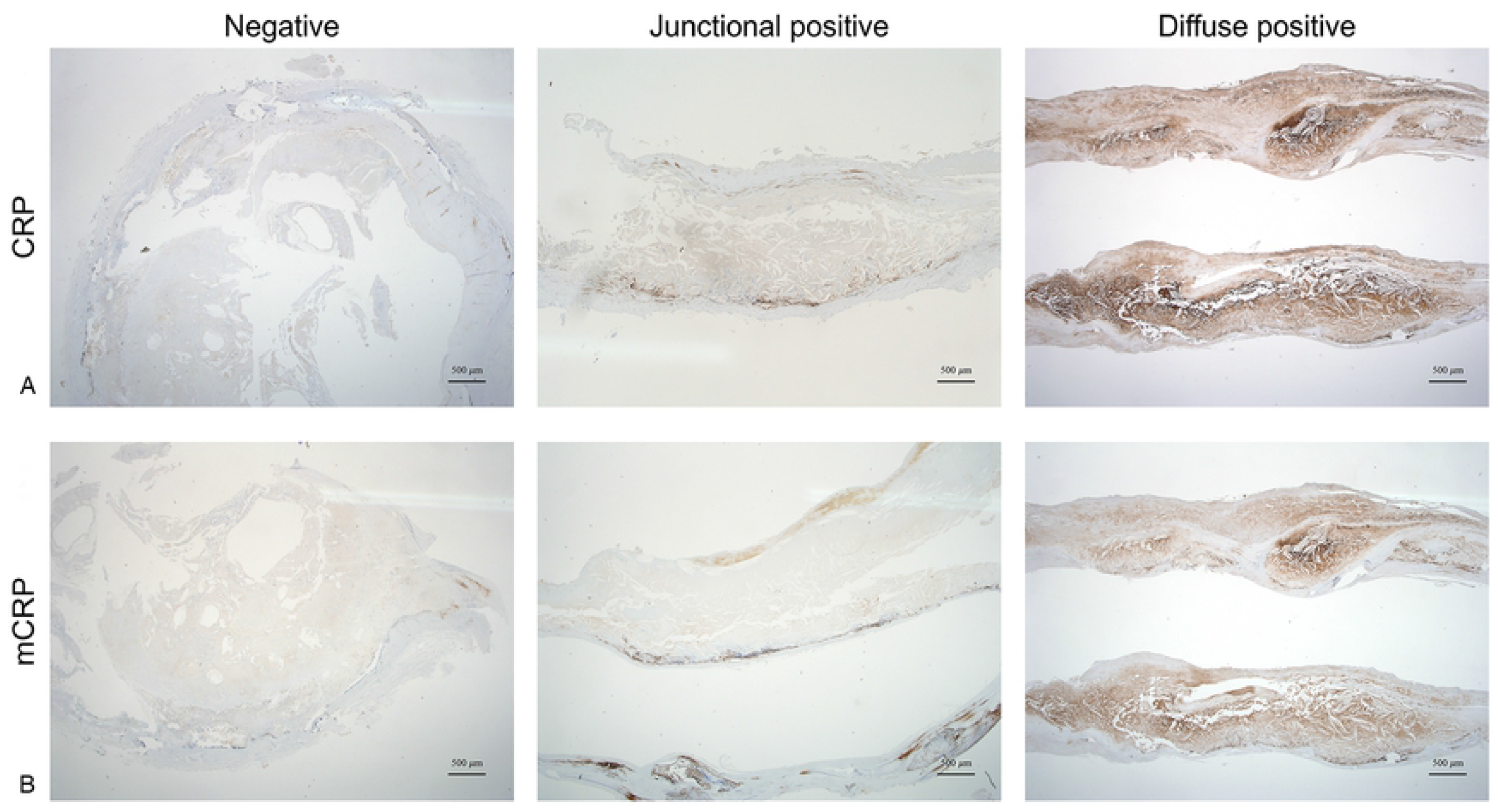
Representative microscopic images showing the patterns of CRP and mCRP immunopositivity in cases of abdominal aortic aneurysm. (A. anti-CRP antibody, B. anti-mCRP antibody, × 12.5).

**Fig 2.**
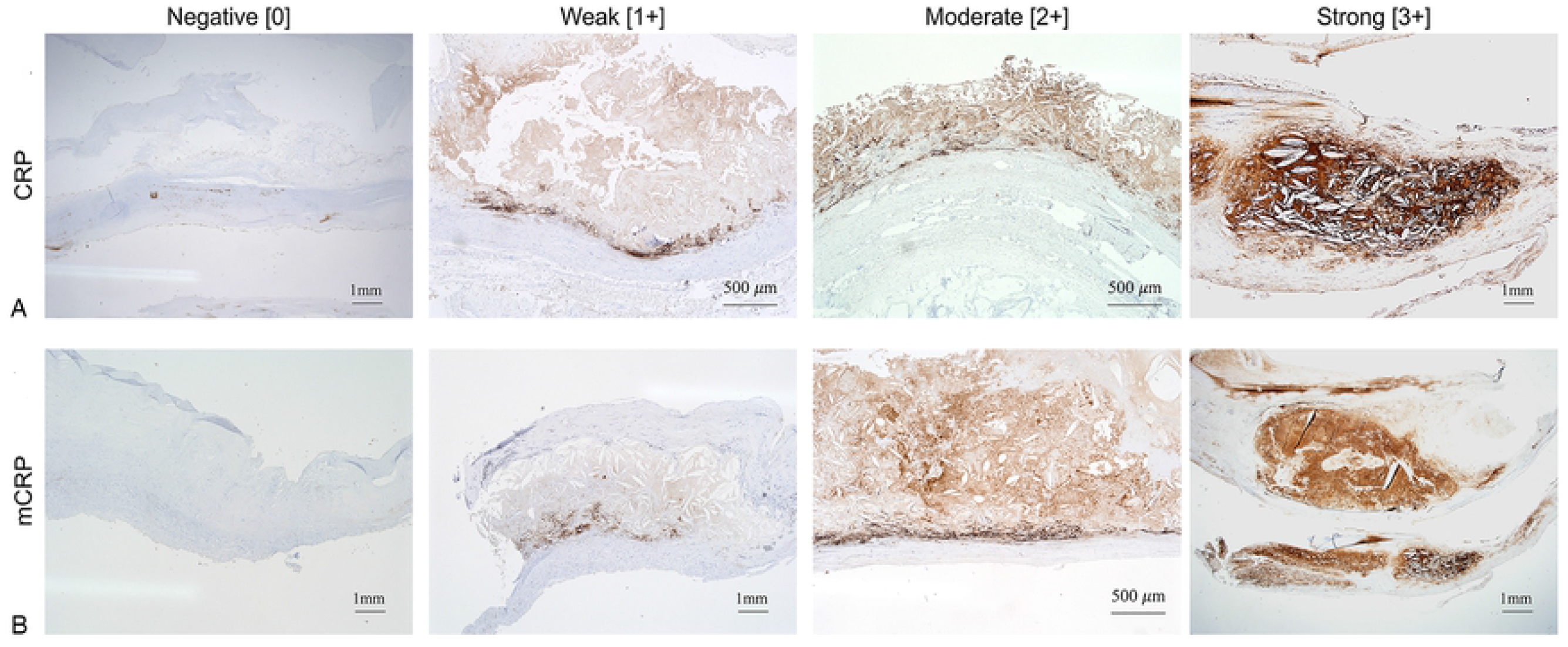
CRP immunostaining scores in the atherosclerosis cores of abdominal aortic aneurysm. A qualitative score between 0 and 3 (negative [0], weak [1+], moderate [2+], strong [3+]) was applied to the immunostaining intensity in each atheroma (A. anti-CRP antibody, B. anti-mCRP antibody).

### Patient selection, and sample preparation for FFPE proteome analysis

By reviewing the slides through H&E and CRP immunostaining, we categorized the patients into (1) AAA-high mCRP (abdominal aortic aneurysm with high [moderate-to-strong (intensity: 2+, 3+)] and diffuse CRP immunopositivity), (2) AAA-low mCRP (abdominal aortic aneurysm with low [negative-to-weak (intensity: 0, 1+)], focal, and junctional CRP immunopositivity), and (3) AAD groups. We randomly selected seven patients from the AAA-high mCRP group, three patients from the AAA-low mCRP group, and two patients from the AAD group for proteomic analysis.

FFPE tissue blocks of the resected aortic specimen were used in this study. Microsections of the samples were meticulously collected under direct microscopic visualization of the H&E-stained slides of FFPE tissue blocks by a pathologist (E.N.K). The atherosclerotic plaque was removed, and only the vessel wall tissues were obtained and used for proteomic analysis. Five sections from unstained slides of dissected materials were collected in 1.5-ml Eppendorf LoBind microcentrifuge tubes (Sigma-Aldrich). Qproteome FFPE tissue kit (Qiagen, Hilden, Germany) was used for deparaffinization and protein extraction, and the protocol was modified to enhance the yield. Briefly, for each sample, 500 μL of heptane was applied in FFPE tissue sections in a 1.5-ml tube and then incubated for 1 hour at room temperature. Subsequently, 25 μL methanol was added to the sample, vigorously vortexed for 10 seconds, and centrifuged at 9000 × g for 2 minutes. The supernatant was discarded, the pellet was dried for 5 minutes. The dried pellet was then dissolved in 94 μL of extraction buffer EXB Plus and 6 μL of ß-mercaptoethanol, and incubated at 4°C for 5 minutes. Next, the samples were incubated in a Thermomixer (Eppendorf AG, Hamburg, Germany) at 100°C for 20 minutes, followed by incubation (750 rpm, 80°C) for 2 hours to break the protein-protein conjugation. After incubation, the tube was placed on ice for 1 minute and then centrifuged for 15 minutes at 14,000 × g at 4°C. The supernatant containing the extracted proteins was transferred to a new tube. Ice-cold acetone was added to each sample to precipitate the proteins, kept at −80°C for 16 hours, and collected by centrifugation at 15,000 × g for 15 min at 4°C. The resulting protein pellet was resuspended in 5% sodium dodecyl sulfate, 50 mM triethylammonium bicarbonate (pH 7.55). Protein concentration was determined by bicinchoninic acid assay.

The extracted proteins were reduced with 20 mM dithiothreitol for 1 hour at room temperature and alkylated using iodoacetamide at a final concentration of 40 mM in the dark for 1 hour. Each sample was loaded onto an S-Trap mini spin column (Protifi, NY, USA) according to the manufacturer’s instructions and digested with trypsin/LysC (1:25 trypsin/protein) for 16 hours at 37°C. The digested peptides were sequentially eluted with i) 50 mM triethylammonium bicarbonate, ii) 0.2% formic acid, iii) 50% acetonitrile/0.2% formic acid. Combined eluates were dried using a speed vacuum and stored at −20°C for liquid chromatography with tandem mass spectrometry (LC-MS) analysis. The overview of the method is depicted in Supplementary Fig S2.

LC-MS analysis was performed as described in our previous analysis[16]. In brief, peptide mixture was trapped and separated on a reversed-phase C18 column (precolumn: Acclaim PepMap, 100 μm × 20 mm, 5 μm, 100 Å, separation column: Acclaim PepMap, 75 μm × 500 mm, 2 μm, 100 Å, Thermo Fisher Scientific) equipped with the Dionex UltiMate 3000 RSLCnano System (Thermo Fisher Scientific, Bremen, Germany). The peptide mixture was separated using gradients from 0% to 40% B buffer at a flow rate of 250 nL/min for 150 minutes with solvent A (5% dimethyl sulfoxide containing 0.1% formic acid) and solvent B (80% acetonitrile containing 5% dimethyl sulfoxide and 0.1% formic acid). Mass spectrum data was collected using Q Exactive Plus mass spectrometer with a nano-ESI source (Thermo Fisher Scientific). A data-dependent mode was applied to acquire mass spectra with a full scan (m/z 350-1800) with 20 data-dependent MS/MS scans. The target number of ions for the full scan MS spectra was 3,000,000, with a maximum injection time of 100 ms and a resolution of 70,000 at m/z 400. The ion target number for MS/MS was set to 1,000,000 with a maximum injection time of 50 ms and a resolution of 17,500 at m/z 400 with normalized collision energy (27%). The dynamic exclusion of repeated peptides was applied for 20 seconds.

The database search was performed as described in our previous study[16]. The acquired spectra were applied on the SequestHT embedded in the Proteome Discoverer (version 2.2, Thermo Fisher Scientific), and the human proteome sequence database (SwissProt database (May 2019)) was used. The precursor mass tolerance was ±10 ppm, and the MS/MS tolerance was 0.02 Da. The default setting of the modification parameters was used, including cysteine carbamidomethylation as a fixed modification and N-terminal/lysine acetylation, methionine oxidation, phospho-serine, phospho-threonine, and phospho-tyrosine as variable modifications with two miscleavages. False discovery rates (FDRs) were set at 1% using “Percolator.” Label-free quantitation was performed using the peak intensity for unique and razor peptides of each protein, and normalization was carried out using the total peptide amount.

The Perseus software (version 1.6.8.0)[17] was used for statistical analysis of the relative abundance of proteins among the samples. The values of normalized protein abundances were transformed into the log2 scale. Three technical replicates of each sample were grouped, and proteins with at least three frequencies among three sample sets were considered as valid values. Missing value imputation of peptides was performed from a normal distribution. Student’s *t*-test was performed using permutation-based FDR (0.01 cut-off) for volcano plots. Hierarchical clustering was performed after z-score normalization. Gene ontology analyses were performed using web-based tools, including the g: Profiler (https://biit.cs.ut.ee/gprofiler/gost) and the Enrichr (https://amp.pharm.mssm.edu/Enrichr/).

Pathway enrichment analyses were performed with the IPA (Qiagen). The enriched canonical pathways and biological functions were based on fold changes and Z-scores. Significantly enriched pathways for the proteins were identified using a cut-off of *P*<0.01. The molecule activity predictor was used to predict the upstream activation or inhibition of a pathway.

C-reactive protein was measured using the Tina-quant C-Reactive Protein Gen.3 reagent (Roche, Basel, Switzerland) designed to achieve very high sensitivity, thereby offering accurate and precise measurements at very low levels of CRP on the Cobas 8000 system (Roche) using the immunoturbidimetric method. Although we did not use the high sensitive (hs)-CRP, the Tina-quant method was shown to be highly associated with hs-CRP[18] and may be suitable for cardiovascular risk assessment[19].

Continuous variables were compared with the Student *t*-test and presented as mean ± SD. Categorical variables were the Fisher’s exact test or Chi-squared test as appropriate and presented as frequencies and percentages. All statistical analyses were performed with the R software version 3.6.2 (R Foundation for Statistical Computing, Vienna, Austria) and SPSS (version 18.0, SPSS Inc., Chicago, USA). All statistical tests were two-sided, and the 5% significance level was used.

## RESULTS

### Clinical characteristics of AAA and AAD

Baseline characteristics of patients with AAA or AAD are shown in Table 1. Risk factors associated with AAA (e.g., male sex, previous cardiovascular disease, smoking, dyslipidemia, arteriosclerosis obliterans) were more prevalent in the AAA group than in the AAD group. Conversely, serum CRP levels and white blood cell count (WBC) were significantly higher in the AAD group than in the AAA group.

**Table 1.** Patient characteristics.

| Characteristics | Abdominal<br>aortic aneurysm<br>(n=69) | Ascending<br>aortic dissection<br>(n=87) | *P* Value |
| --- | --- | --- | --- |
| Age, y | 68.0±8.4 | 62.0±14.9 | 0.002 |
| Male, n (%) | 64 (92.8) | 42 (48.3) | <0.001 |
| Body mass index, kg/m<sup>2</sup> | 24.2±2.8 | 24.7±4.1 | 0.424 |
| Serum CRP level, mg/dL | 0.4±0.6 | 2.7±4.5 | <0.001 |
| WBC, count/μL | 6861±1494 | 11159±4530 | <0.001 |
| Diabetes mellitus, n (%) | 9 (13.0) | 4 (4.6) | 0.109 |
| Hypertension, n (%) | 50 (72.5) | 80 (92.0) | 0.002 |
| Previous cardiovascular disease<sup>a</sup>, n (%) | 21 (30.4) | 9 (10.3) | 0.003 |
| Alcohol consumption, n (%) | 36 (52.2) | 41 (47.1) | 0.642 |
| Smoking, n (%) | 48 (69.6) | 29 (33.3) | <0.001 |
| Dyslipidemia<sup>b</sup>, n (%) | 44 (63.8) | 35 (40.2) | 0.006 |
| Arteriosclerosis obliterans, n (%) | 35 (50.7) | 1 (1.1) | <0.001 |
| Tuberculosis, n (%) | 5 (7.2) | 3 (3.4) | 0.482 |

<sup>a</sup> Included history of stroke, myocardial infarction, or angina requiring percutaneous coronary intervention
<sup>b</sup> Patients who were on statin medication.

### Histologic analysis

In all cases of AAA, atheromas were observed in the lumen of the aneurysmal wall. Thinning of the aortic wall media and diminishing of the elastic lamella that was in contact with the atheroma with fibrous cap were observed as well. In the AAD cases, as we selected non-atheromatous aortic dissection specimens, atheromas were not observed, and thinning of the aortic walls was not observed; instead, all cases of AAD showed medial tearing (Supplementary Fig S3).

We observed that 95.8% (23/24) of AAA cases showed strong immunopositivity of both CRP and mCRP in the aneurysmal aortic wall. In particular, CRP immunopositivity was evident in the junction between atherosclerotic plaque and eroded aortic media, which is where the elastic lamina was diminished (Fig 3 A–D). On the other hand, both CRP and mCRP were not immunopositive in any of the AAD cases (Fig 3 E–H).

**Fig 3.**
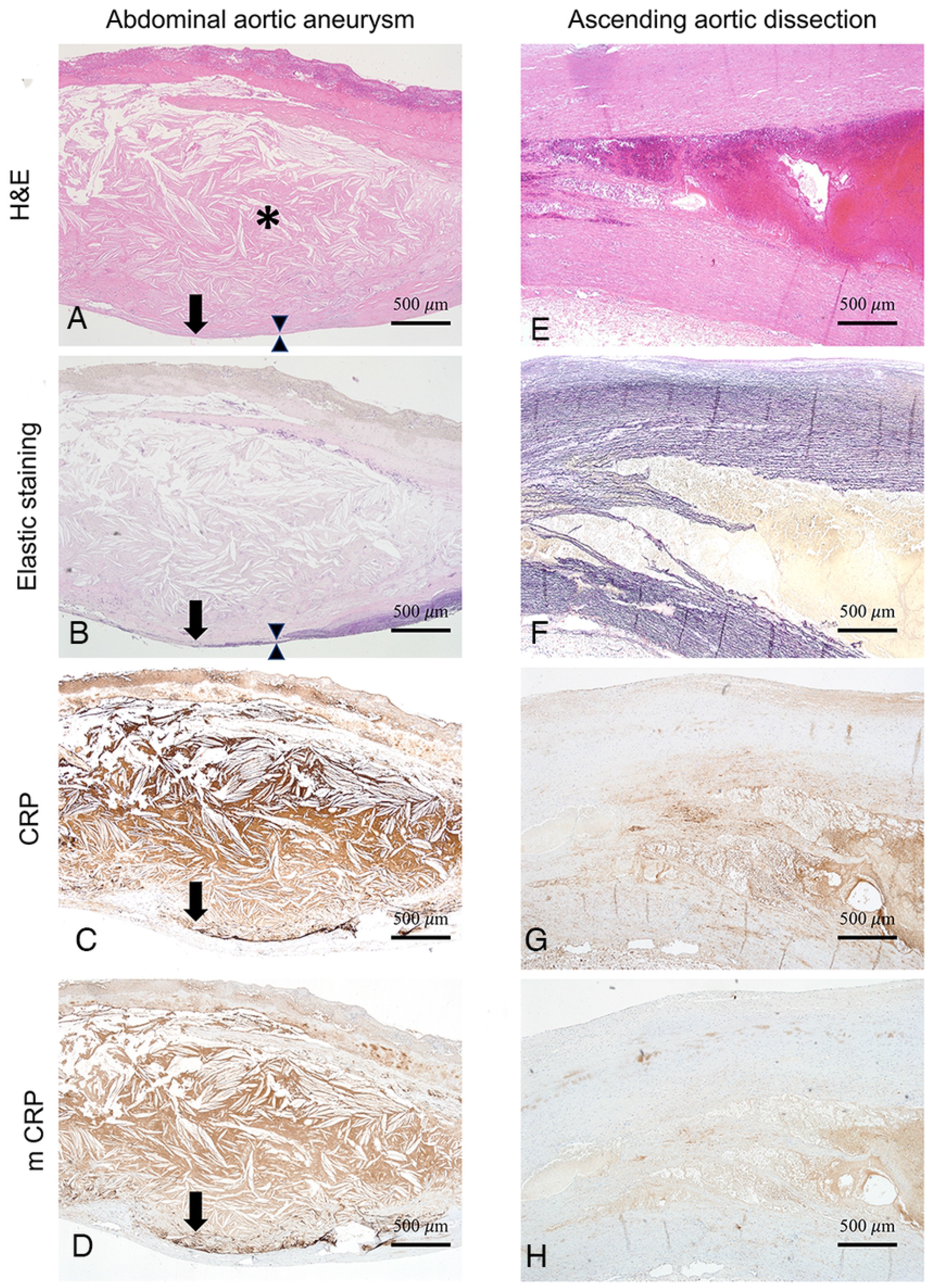
CRP immunopositivity with the corresponding histopathologic findings. (A–D) Aorta specimen from a patient with abdominal aortic aneurysm and mildly elevated serum CRP (0.28 mg/dL). The site of strong and linear CRP (C) and mCRP (D) immunopositivity was found in the interface (arrow) between atherosclerotic plaque (asterisk) and the thinned aneurysmal wall (between the two arrowheads). This immunopositive lesion was correlated with the area of the diminished elastic lamella of the eroded media (B). Atheroma was diffusely immunopositive for both CRP and mCRP. (A, H&E, B, elastic staining, C, anti-CRP antibody, D, anti-mCRP antibody, × 40). (E–H) Aorta specimen from a patient with ascending aortic dissection with mildly elevated serum CRP (0.74 mg/dL). CRP and mCRP were faintly and nonspecifically stained in the smooth muscle cell of tunica media. CRP was not stained at the boundary of the damaged and the torn aortic wall. (E, H&E, F, elastic staining, G, anti-CRP antibody, H, anti-mCRP antibody, × 40).

In terms of the distribution of anti-CRP immunopositivity, 67% (16/24) and 29% (7/24) of AAA cases showed junctional and diffuse immunopositivity, respectively. mCRP immunopositivity showed a similar pattern, with 71% (17/24) and 25% (6/24) of AAA cases showing junctional and diffuse immunopositivity, respectively (Fig 1). For CRP, the immunopositivity was categorized as negative [0], weak [1+], moderate [2+], and strong [3+] in 4.2% (1/24), 50% (12/24), 16.7% (4/24), and 29.2% (7/24) of the AAA cases, respectively, and 4.2% (1/24), 54% (13/24), 21% (5/24), and 21% (5/24) for mCRP, respectively (Fig 2).

Serum CRP >0.1 mg/dL was associated with stronger intensity and the larger area of CRP immunopositivity within atheromas and the aortic wall (Fig 4, 5). In patients with low serum CRP (≤0.1 mg/dL), CRP and mCRP immunopositivity were weak in 64.3% and 78.6%, respectively; in contrast, among patients with elevated serum CRP (>0.1 mg/dL), CRP and mCRP showed moderate-to-strong immunopositivity in 70.0% and 80.0% of the cases (Supplementary Table S1); such trend of immunopositivity strengths was significantly different between the two groups (CRP, *P*=0.002; mCRP, *P*=0.007). Most of the patients (92.9%) with low serum CRP showed junctional immunopositivity and none showed diffuse immunopositivity for both anti-CRP and mCRP, whereas those with elevated serum CRP showed junctional positivity only in 30.0% and 40.0% and diffuse positivity in as much as 70.0% and 60.0% for CRP and mCRP, respectively; such difference in the staining patterns was statistically significant (CRP, *P*=0.001; mCRP, *P*=0.003).

**Fig 4.**
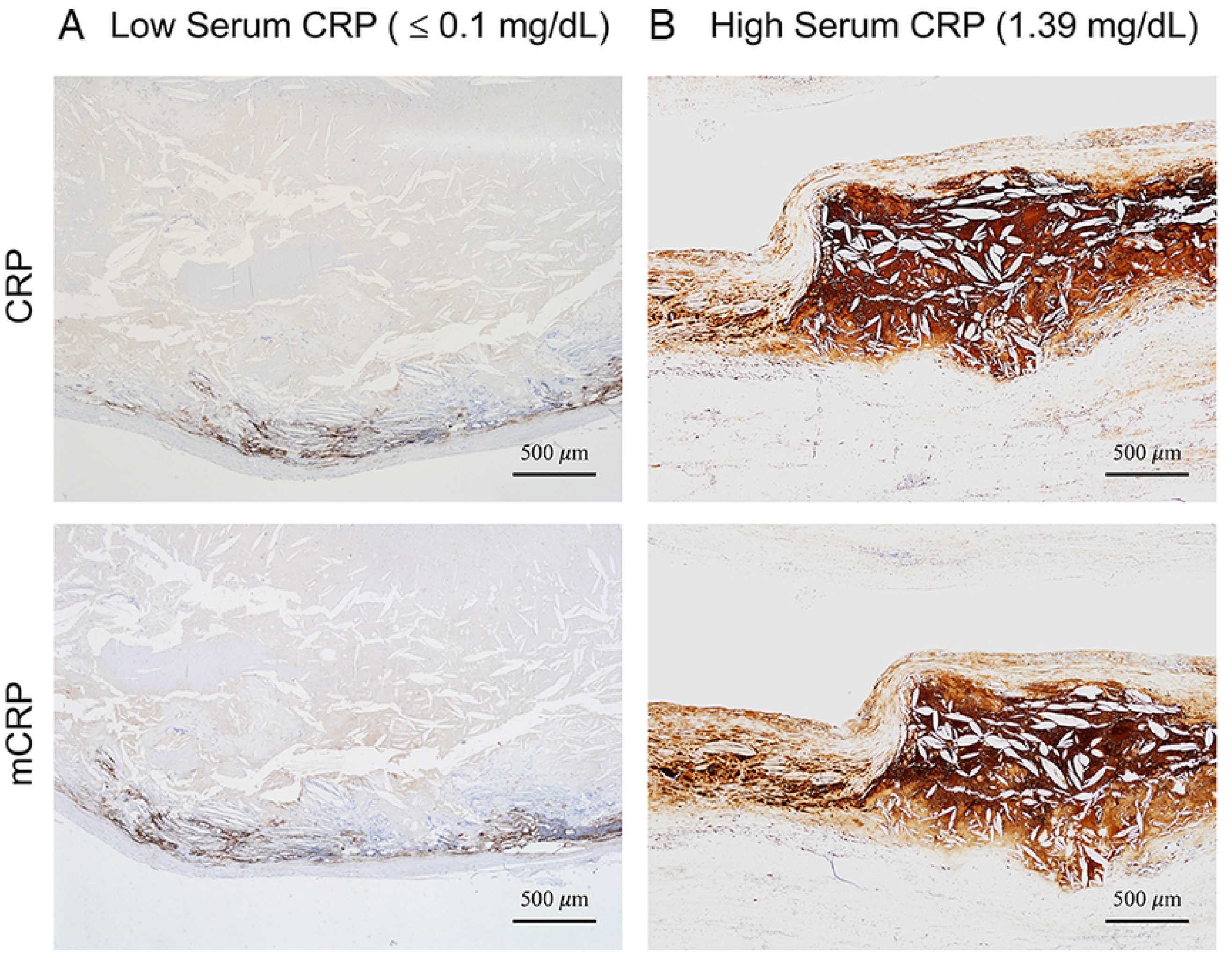
Representative CRP and mCRP immunostaining according to serum CRP levels. (A) Low serum CRP (≤0.1 mg/dL). (B) High serum CRP (1.39 mg/dL) (anti-CRP antibody, anti-mCRP antibody, × 40).

**Fig 5.**
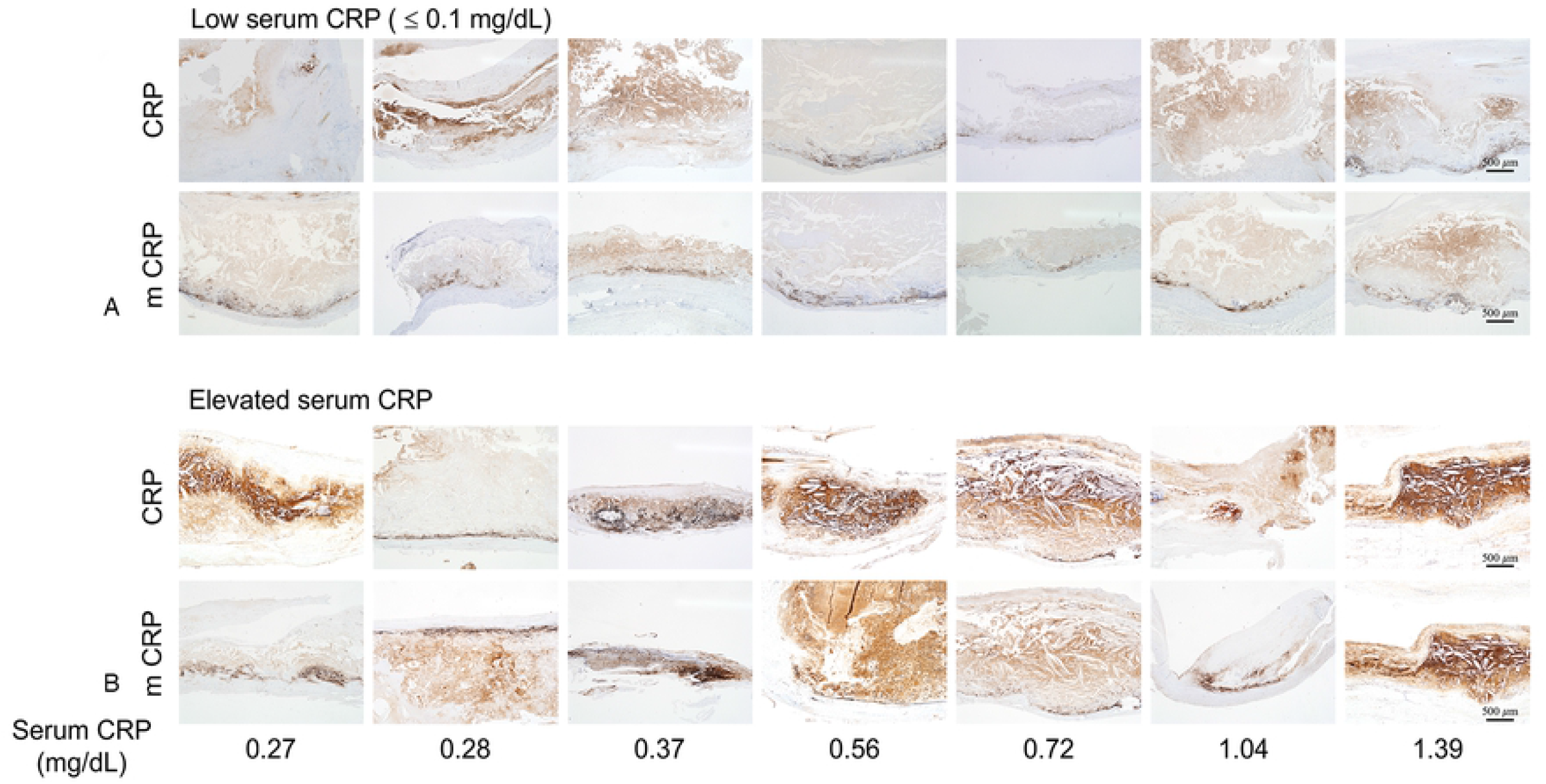
CRP and mCRP immunostaining according to serum CRP level (a, ≤0.1 mg/dL; b, >0.1 mg/dL) in cases of abdominal aortic aneurysm with atherosclerosis (all magnification × 40).

Because the CRP immunostaining pattern was significantly different according to CRP cut-off level of 0.1 mg/dL, we divided the AAA cases into the low serum CRP group (≤ 0.1 mg/dL; n=31) and the elevated serum CRP group (>0.1 mg/dL; n=38) and analyzed the patient characteristics (Supplementary Table S2); as a result, we found that the maximal diameter of aortic aneurysm was significantly larger in the elevated serum CRP group (6.5 ± 1.5 cm) than in the low serum CRP group (5.7 ± 1.2 cm, *P*=0.013). The elevated serum CRP group also showed strong immunopositivity for MCP-1, C3a, and C5a along with CD68-positive macrophages within the atheromas and degenerated aortic walls (Fig 6).

**Fig 6.**
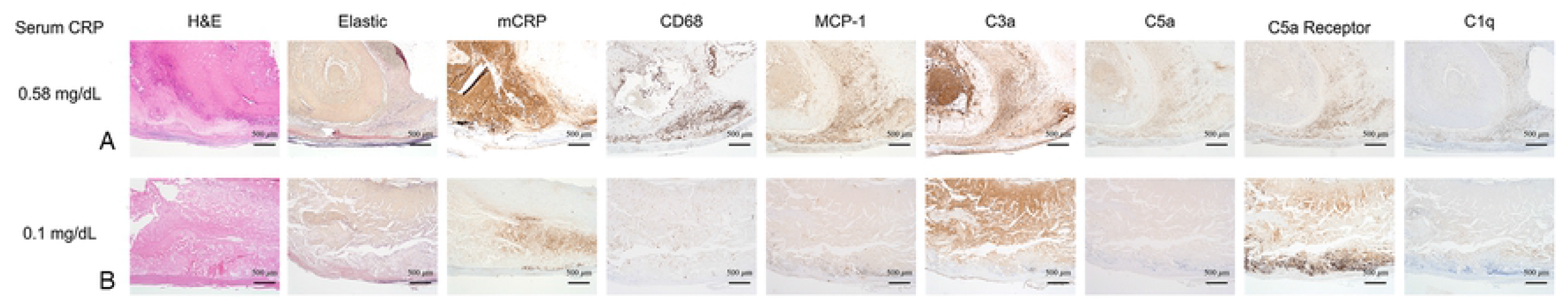
Representative images of aortic specimens from cases of AAA according to serum CRP levels. H&E, elastic staining, and the immunostaining patterns of mCRP, CD68, MCP-1, and complement components (C3a, C5a, C5a receptor, C1q) in patients with high serum CRP level (0.58 mg/dL, a) and low serum CRP level (0.1 mg/dL, b). (× 40; MCP-1, monocyte chemoattractant protein-1)

### LC-MS proteomics analysis

By incorporating the results of CRP immunostaining, the study groups for proteomic analysis were divided into AAA with strong and diffuse CRP immunopositivity (AAA-high mCRP, n=7), AAA with weak, focal and junctional CRP immunopositivity (AAA-low mCRP, n=3), and AAD (n=2) groups. The demographics of the three groups are described in Supplementary Table S3.

We carried out three separate LC-MS/MS analyses for each aortic sample for protein profiling. After normalization, the triplicate results were analyzed for protein abundance. The protein abundance in the triplicate samples was consistent, confirming that protein quantification was of high quality. Pearson correlation analysis across the samples showed a high correlation among triplicates, thus indicating good experimental reproducibility (R = 0.93, Supplementary Fig S4). Correlation tests with a heatmap in each group using quantitative protein information obtained through MS analysis showed a high correlation among the three groups, indicating reasonable experimental reproducibility (Fig 7A). Two-dimensional principal component analysis (PCA) plots using quantitative protein information indicated that the three groups were sufficiently distinct and that each group had different characteristics in quantitative protein profiles (Fig 7B).

**Fig 7.**
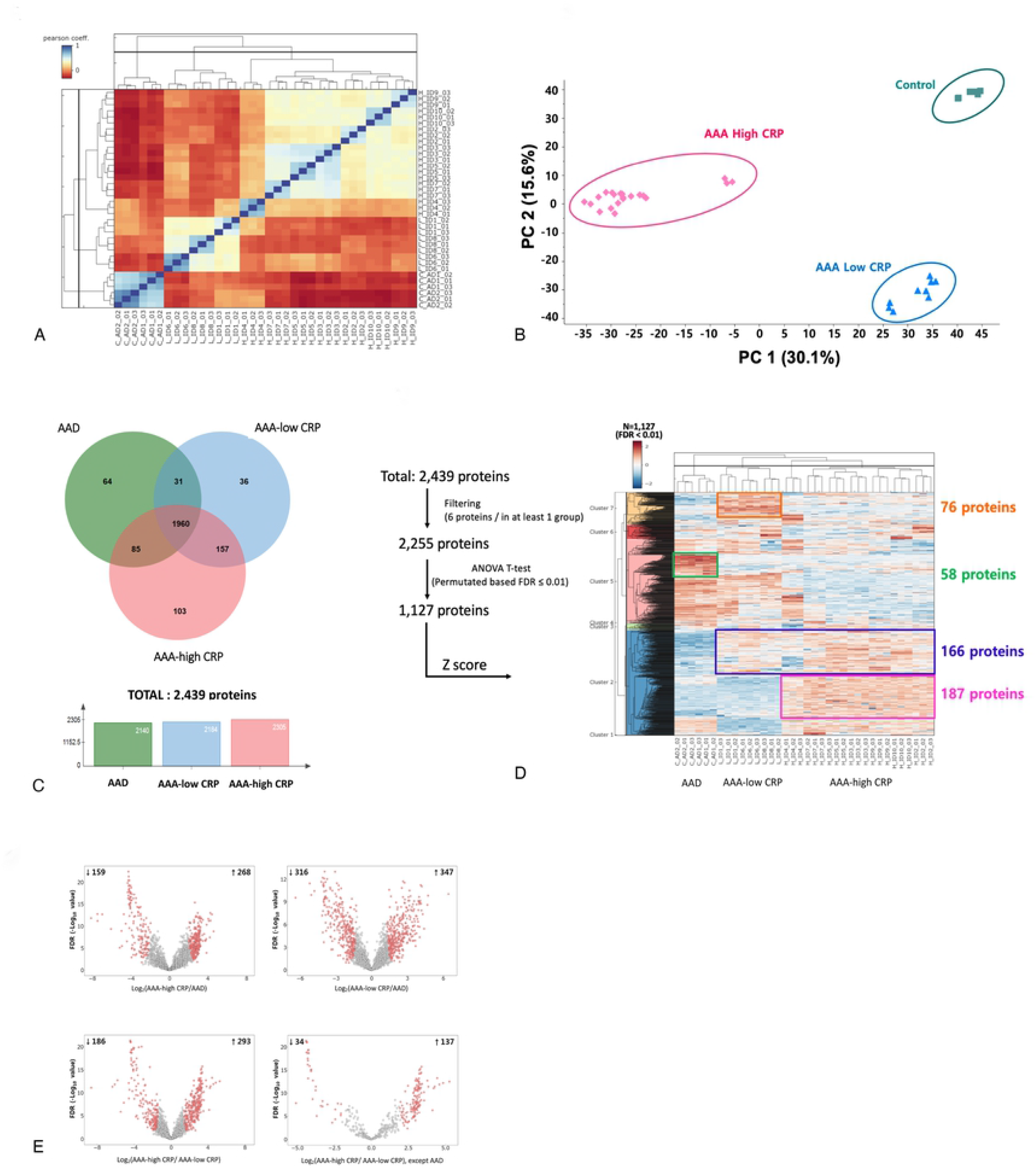
Proteomic profiles and comparison of identified proteomes among the AAA-high mCRP, AAA-low mCRP, and AAD groups. (A) Correlation test with a heatmap in each group using quantitative protein information obtained through MS analysis. (B) Two-dimensional principal component analysis plots using quantitative protein information. (C) Venn diagram showing the overlap of differentially expressed proteins among the three groups. (D) Hierarchical clustering of top abundant 1,127 DEPs between AAD, AAA-low mCRP, and AAA-high mCRP groups (ANOVA *t*-test, permutation-based FDR ≤0.01). Rows represent proteins and columns represent different samples. Darker shades of red and blue each indicate increased and decreased expressions compared with control. This figure was generated using Instant Clue (version 0.9.2 from http://www.instantclue.uni-koeln.de/) (E) Volcano plots illustrating significantly differentially abundant proteins between AAD, AAA-low mCRP, and AAA-high mCRP groups. Each dot indicates a protein and red colors indicate significantly enriched proteins with *q* values <0.05 and fold changes of more than ± 2-fold change, and permutation-base FDR ≤0.01. AAA-low mCRP indicates aortic aneurysm with weak and focal CRP immunopositivity; AAA-high mCRP, Aortic aneurysm with strong and diffuse CRP; AAD, ascending aortic dissection.

A total of 2,439 proteins were identified in the three groups—2305 in AAA-high mCRP, 2184 in AAA-low mCRP, and 2140 in AAD. Venn diagram analysis showed that 103, 36, and 64 proteins were exclusively identified in the AAA-high mCRP, AAA-low mCRP, and AAD groups, respectively (Fig 7C). For further analysis, the 2,439 proteins were filtered by ANOVA in the three groups with permutation-based FDR ≤ 0.01, and 1,127 proteins were selected. Subsequent hierarchical clustering analysis revealed 58 proteins that were differentially expressed only in the AAD group, 187 proteins were differentially expressed only in the AAA-high mCRP group, and 166 proteins that were commonly expressed in the AAA-high mCRP and AAA-low mCRP groups and not in the AAD group. A heatmap of clustered fold changes for the DEPs is shown in Fig 7D.

Compared with the AAD group, AAA-high mCRP and AAA-low mCRP groups showed differential abundance in 427 and 663 proteins, respectively, and 479 proteins were differentially abundant between AAA-high mCRP and AAA-low mCRP groups. Excluding the DEPs of the AAD group, 171 proteins were differentially expressed between the AAA-high mCRP group and the AAA-low mCRP group (fold change >2 or ≤0.5, FDR<0.01, Supplementary Table S4). The list of the 171 DEPs is provided in Supplementary Dataset online. Of the DEPs, the 11 top-ranking proteins were chosen by combining the protein abundance ratio and the Student’s t-test *P* values (Supplementary Table S5), including glutamine-dependent NAD (+) synthetase, glucose 14-3-3 protein sigma, glucosamine-6-phosphate isomerase 1, involucrin, and peptidyl-prolyl cis-trans isomerase FK506-binding protein 2 (FKBP2). Fig 7E shows the volcano plots of proteins that were differentially abundant among groups.

Enrichr analysis revealed that 187 proteins were more abundant in both AAA-high mCRP and AAA-low mCRP than in AAD. The entries of the 187 proteins were analyzed according to biological processes, molecular function, and cellular component (Fig 8A). The top enriched biological processes in the AAA-high mCRP group were regulation of protein activation cascade, regulation of complement activation, and regulation of humoral immune response. The top enriched biological processes in the AAA-low mCRP were neutrophil regulation, neutrophil activation involved in immune response, and neutrophil-mediated immunity. Interestingly, ficolin-1-rich granule cellular component was specifically enriched in the AAA-low mCRP group.

**Fig 8.**
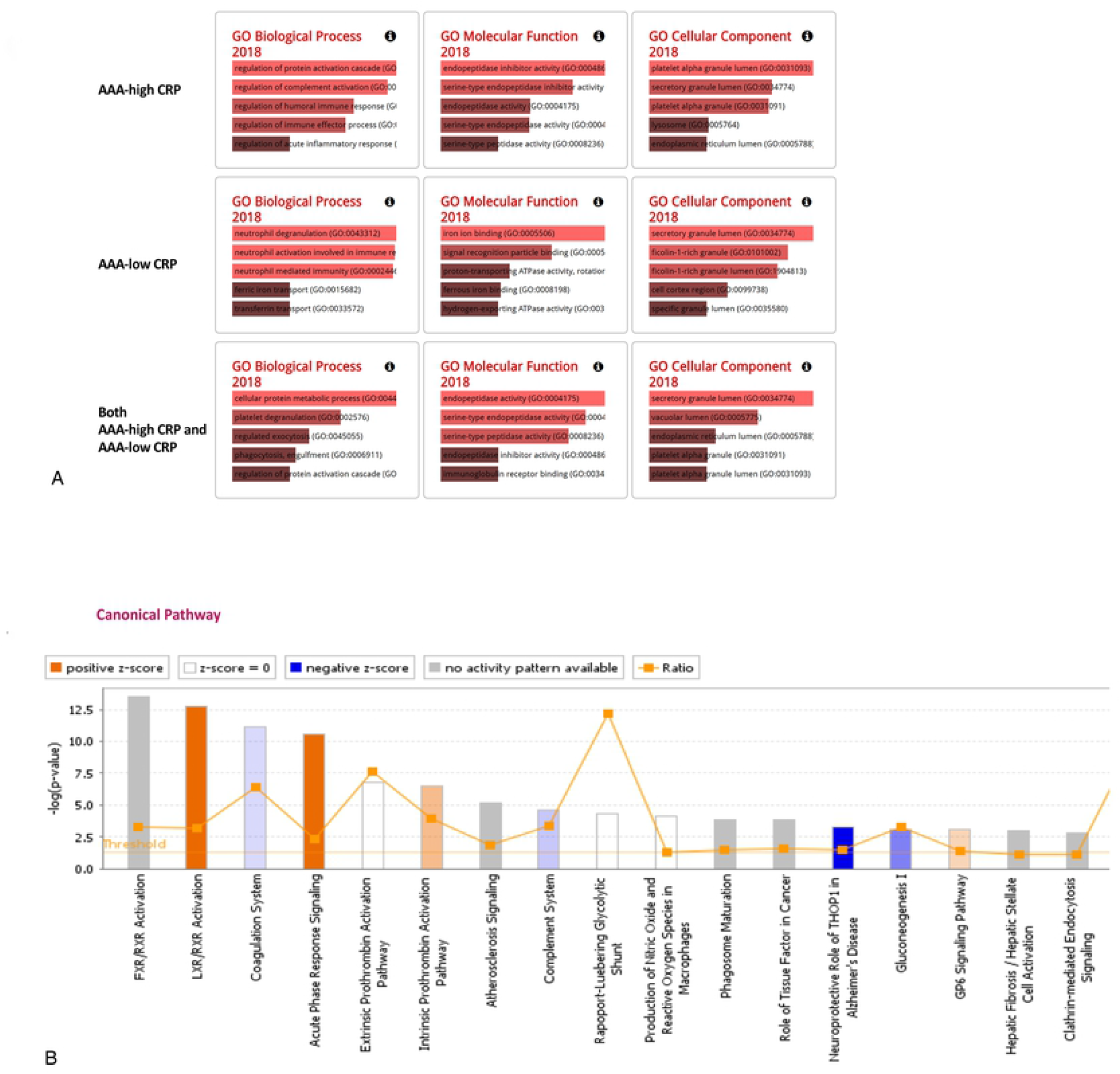
Bioinformatics analysis of 187 proteins that showed higher abundance in both AAA-high mCRP and AAA-low mCRP compared with AAD. (A) Gene ontology analysis. Top 5 subcategories in biologic process, molecular function, and cellular component. (B) Top canonical pathways identified in core analysis in IPA. The bar graph shows the canonical pathway represented by gene enrichment. The orange line running through the bars shows the threshold for the *P*-value for the particular pathway’s enrichment. The vertical axis is the −log(p-value) and the horizontal axis shows the given pathways.

IPA (Qiagen) was performed on 171 proteins whose expressions were significantly different between the AAA-high mCRP group and the AAA-low mCRP group after excluding the DEPs of the AAD group. Multiple canonical signaling pathways including liver X receptor/retinoid X receptor (LXR/RXR) activation, coagulation system, acute phase response signaling, extrinsic and intrinsic prothrombin activation pathway, atherosclerosis signaling, and complement systems were significantly more enriched in the AAA-high mCRP group (Fig 8B, 9). Notably, the pathways in acute phase response signaling included those resulting in the production of CRP (Fig 9C) along with C3 and C4. In Disease or Functions Annotation analysis, multiple proteins related to engulfment of cells, phagocytes, interaction of leukocytes, migration of phagocytes, and response granulocytes were predicted as significantly more activated in the AAA-high mCRP group than in the AAA-low mCRP group (Fig 9H).

**Fig 9.**
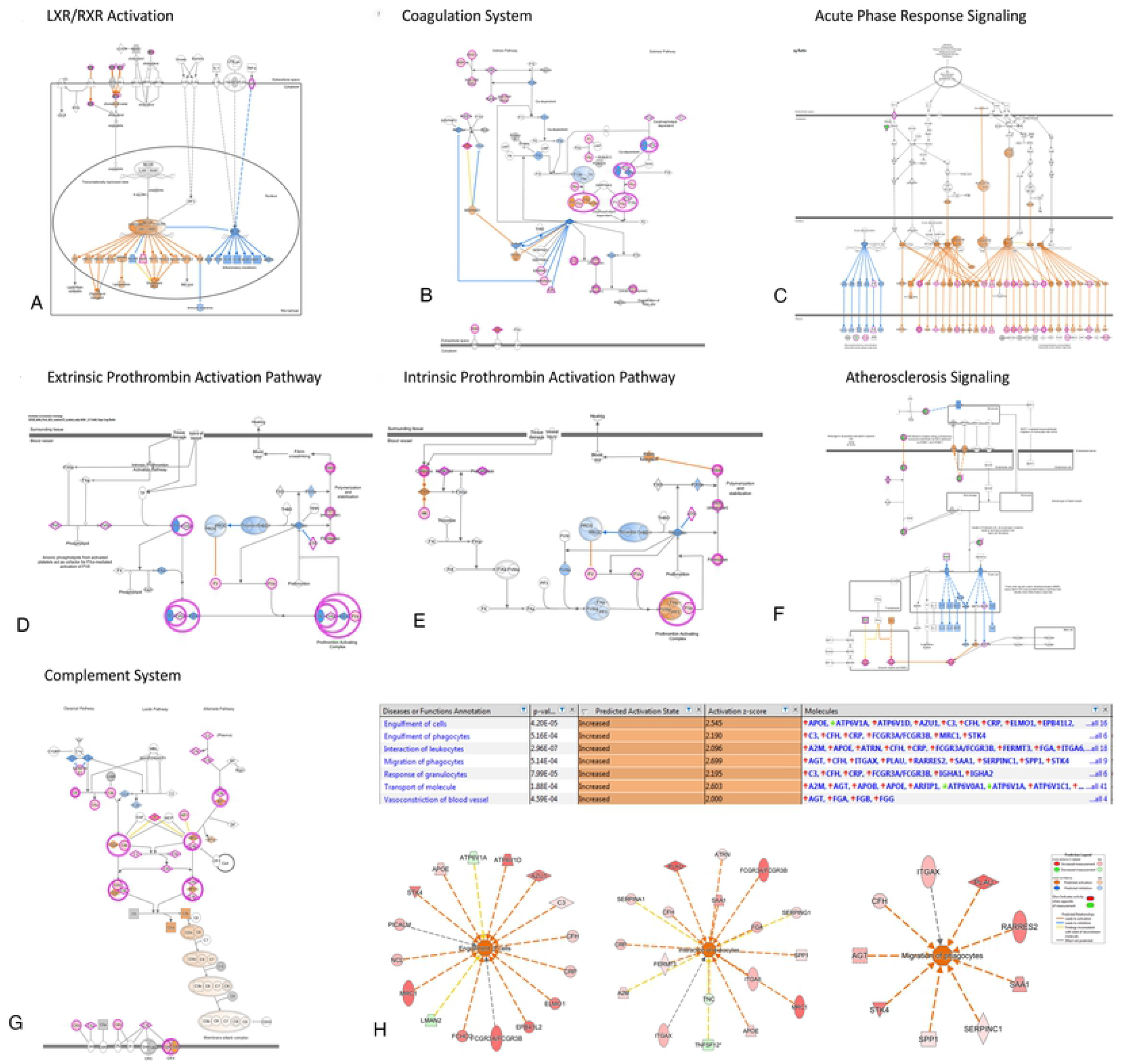
Canonical pathway analysis of signaling pathways. IPA bio function analysis (QIAGEN Inc., https://www.qiagenbioinformatics.com/products/ingenuity-pathway-analysis/) was carried out to assemble a network between (A) LXR/RXR Activation, (B) coagulation system, (C) acute phase response signaling, (D) extrinsic prothrombin activation pathway, (E) intrinsic prothrombin activation pathway, (F) atherosclerosis signaling, and (G) complement system of the proteins differentially abundant in the AAA-high mCRP group. Red and green colors represent the measured levels of increased and decreased molecules, respectively. Orange and blue colors represent the predicted activation or inhibition of molecules or bio functions, respectively. (H) Annotation enrichment analysis. Networks including “engulfment of cells,” “the interaction of leukocytes,” and “migration of phagocytes” were significantly enriched in the AAA-high mCRP group than in the AAA-low mCRP group.

## DISCUSSION

We found that mCRP was deposited in the lumen of dilated aneurysmal sacs that are in contact with the atheroma, and that internal elastic lamellae had disappeared at the site of mCRP deposition. In AAD, mCRP was not deposited in the dissected aortic wall despite elevated serum CRP level. By using proteomic analysis, we found that various pathologic signaling pathways such as LXR/RXR activation, atherosclerosis, and complement activation were upregulated in the mCRP-deposited dilated aortic walls with high serum CRP level.

We observed a correlation between serum CRP level and aortic aneurysmal sac size. Previous studies showed that serum CRP levels were associated with large aortic aneurysm diameters[20]. Also, we found that patients with elevated serum CRP levels had strong, diffuse patterns of both anti-CRP and anti-mCRP immunostaining in the aortic aneurysmal walls. Accordingly, this observation was confirmed by proteome analysis in which pathologic signaling pathways associated with mCRP deposition were upregulated in AAA cases with high CRP levels. In its native state, CRP exists in a pentameric form in the serum; however, upon contact with damaged cells, CRP dissociates and transforms into a monomeric form and promotes inflammation to exacerbate tissue damage[10]. We suggest that pCRPs secreted from aneurysmal sacs and atheroma come into contact with the degenerated cell membrane of the aortic walls in AAA, deposited as mCRPs along with the increase of serum CRP, and accelerate tissue damage.

Inflammation has a major role in the pathogenesis of AAA through the production of proteolytic enzymes, oxidation-derived free radicals, and cytokines[3]. We observed strong immunopositivity of MCP-1 in strong mCRP-deposited AAA; moreover, IPA showed that the LXR/RXR activation pathway was highly enriched in the AAA with high mCRP group, which is a principal pathway involved in the regulation of inflammation[21], lipid metabolism[22], and atherogenesis[23]. The proteomic profiling of mCRP-deposited AAA tissues with high serum CRP levels showed marked upregulations of complement activation, humoral immune response, acute inflammatory response, engulfment, migration of phagocytes, interaction of leukocytes, and response of granulocytes. These results well-support the inflammation hypothesis of AAA development and the role of mCRP deposition and serum CRP elevation thereof[24].

Interestingly, we observed that mCRP was specifically deposited in the interface between the thinned aortic walls and atherosclerotic plaque in AAA, a region that corresponded to the site of the degenerated elastic lamina. The proteomic analysis also showed that the pathway associated with atherosclerosis was activated in the AAA-high mCRP group (Fig 9F). The role of atherosclerosis in the pathogenesis of AAA remains controversial[3]; however, most cases of AAA are regarded to be associated with atherosclerosis because AAA is mostly observed above or with atherosclerosis[25] and because the atherosclerotic plaque on the aortic wall is associated with the erosion of the aortic media[26]. Our finding that the site of mCRP deposition was between atherosclerosis and the degenerated aortic wall suggests that there is a cross-talk between CRP and atherosclerosis in the pathogenesis of AAA.

In patients with elevated serum CRP levels and strong and diffuse positivity of mCRP, MCP-1 and complement components were also immunopositive in the atheroma and CD68^+^ macrophages in AAA (Fig 6A). Global pathway analysis revealed several upregulated proteins related to complement activation or acute phase response, including CRP (Fig 9C). The deposition of mCRP in damaged tissue induces complement binding[27], and CRP leads to more C1q binding and C1 activation in its monomeric form than the pentameric form[28]. The deposition of mCRP also increases the concentration of inflammatory chemokines[29]. In terms of aortic aneurysm, complement activation contributes to the progression of AAA[30]. Collectively, the interactions among mCRP deposition, chemokine, and complement components may act as a mechanism for the exacerbation of AAA.

Elastin is a crucial structural component of the aorta, and its degradation by proteolytic enzymes such as matrix metalloproteinases (MMP) contributes to the development of aortic aneurysm[31]. Our IPA results showed that MMP-1 was increased in the AAA-high mCRP group during the activation of the atherosclerosis pathway (Fig 9F). Thus, we suggest that the deposition of mCRP on the aortic wall of the aneurysms may damage the aortic wall by inducing atherosclerosis through MMP-1. Accordingly, CRP has been shown to upregulate MMPs in acute myocardial infarction[32] while being associated with plaque rupture[33].

In our study, the coagulation systems of both intrinsic and extrinsic prothrombin activation pathways were enriched in the AAA-high mCRP group. Intraluminal thrombus stimulates AAA progression by inducing localized hypoxia at the underlying aortic wall[34], which in turn triggers adventitial angiogenesis and aggravates inflammatory infiltration from the outer vessel layers[35]. Intraluminal thrombus thickness is correlated with AAA diameter, elastin degradation, and smooth muscle cell apoptosis[36], and proteomic analysis of intraluminal thrombus showed that complements are activated in AAA[37]. Moreover, thrombus in AAA entraps neutrophils to create a pro-oxidant and proteolytic environment that leads to the aggravation of aortic aneurysm[38]. Therefore, we suggest that mCRP deposition may aggravate AAA through thrombus formation.

The results of our gene ontology analysis showed that mFicolin-1-rich granules were enriched in the AAA-low mCRP group compared with the AAA-high mCRP group. Secreted mFicolin has a protective effect by anchoring onto monocyte transmembrane G protein-coupled receptor 43 and crosstalks with CRP to curtail the production of IL-8, thereby preventing immune overactivation[39]. Considering that the AAA-high mCRP group was relatively vulnerable to immune reaction compared with the AAA-low mCRP group, we suggest that mFicolin could have acted as a protective factor of aortic aneurysm.

Interestingly, despite the significantly elevated serum CRP level, AAD did not show depositions of mCRP. The presence of atherosclerosis was one of the most important differences between the AAA and AAD groups, and considering that the site of mCRP deposition was where atherosclerosis and the aortic wall were in contact, we suggest that the presence of atherosclerosis is an essential factor for the deposition of mCRP.

Several limitations of this study should be acknowledged. First, for comparison with AAA, we used tissue specimens from patients with AAD rather than healthy individuals with normal aortas. As such, the possibility of differences in protein expression arising from the difference in the anatomical location of the tissues or the disease characteristics cannot be ruled out. Alternatively, aorta specimens obtained from heart transplantation could have been used in place of the normal aorta. However, we deemed that such specimens would harbor other confounding factors arising from the respective underlying cardiovascular conditions and thus be unsuitable for use as a control. Therefore, we decided to use AAD, which is largely associated with hypertension than atherosclerosis[40]. Second, considering the small number of study patients, especially those used in the proteomic analysis, our results should be mainly considered as hypothesis-generating. In order to gather a homogenous group of cases with AAA or AAD, we excluded many aortic diseases such as connective or genetic aortic diseases. Nevertheless, each group showed high intragroup correlation in the LC-MS analysis, indicating that the pathophysiology and signaling pathways were distinctly grouped even with a small number of patients. Third, the causality between mCRP deposition and AAA aggravation could not be determined due to the retrospective nature of the study. Lastly, we could not discriminate whether proteome change pattern is due to the deposition of mCRP itself or underlying serum CRP. Further mechanistic studies will provide confirmatory data on the pathogenic mechanisms in AAA regarding mCRP deposition[3].

## CONCLUSION

In conclusion, our results show that mCRP is specifically deposited on the interface between atheroma and the damaged aortic wall in AAA. Proteomics analysis revealed that deposition of mCRP in the aneurysmal wall along with high serum CRP level is associated with various signaling pathways related to complement activation, atherosclerosis, and thrombogenesis. These results collectively suggest that mCRP deposition accompanied by increased serum CRP has a possible role in the pathological process of aortic aneurysm, and provide novel insight for halting the progression of the aortic aneurysm by the discovery of druggable protein targets.

## Abbreviations

AAA: abdominal aortic aneurysm
AAA-high mCRP: abdominal aortic aneurysm with strong and diffuse mCRP immunopositivity
AAA-low mCRP: abdominal aortic aneurysm with weak, focal and junctional mCRP immunopositivity
AAD: ascending aortic dissection
CRP: C-reactive protein
DEPs: differentially expressed proteins
FDR: false discovery rate
FFPE: formalin-fixed paraffin-embedded
FKBP2: FK506-binding protein 2
H&E: hematoxylin and eosin
hsCRP: high sensitive C-reactive protein
IPA: ingenuity pathway analysis
IRB: institutional review board
LC-MS/MS: liquid chromatography with tandem mass spectrometry
LXR/RXR: liver X receptor/retinoid X receptor
MCP-1: monocyte chemoattractant protein-1
mCRP: monomeric form of C-reactive protein
MMPs: matrix metalloproteinases
PCA: principal component analysis
pCRP: pentameric form of C-reactive protein
WBC: white blood cell count

## Supporting information

### Supplementary Dataset online

The list of the 171 differentially expressed proteins between the AAA-high mCRP group and the AAA-low mCRP group

